# Defining cellular population dynamics at single cell resolution during prostate cancer progression

**DOI:** 10.1101/2022.03.02.482711

**Authors:** Alexandre A. Germanos, Sonali Arora, Ye Zheng, Erica T. Goddard, Ilsa M. Coleman, Anson T. Ku, Scott Wilkinson, Robert A. Amezquita, Michael Zager, Annalysa Long, Yu Chi Yang, Jason H. Bielas, Raphael Gottardo, Cyrus M. Ghajar, Peter S. Nelson, Adam G. Sowalsky, Manu Setty, Andrew C. Hsieh

**Affiliations:** Division of Human Biology, Fred Hutchinson Cancer Research Center, Seattle, WA 98109; University of Washington Molecular and Cellular Biology Program, Seattle, WA 98195; Division of Vaccine and infectious Diseases, Fred Hutchinson Cancer Research Center, Seattle, WA 98109; Division of Public Health Sciences, Translational Research Program, Fred Hutchinson Cancer Research Center, Seattle, WA 98109; Laboratory of Genitourinary Cancer Pathogenesis, Center for Cancer Research, National Cancer Institute, NIH, Bethesda, MD 20892; Center for Data Visualization, Fred Hutchinson Cancer Research Center, Seattle, WA 98109; University of Washington Departments of Medicine and Genome Sciences, Seattle, WA 98109; Translational Data Science Integrated Research Center, Fred Hutchinson Cancer Research Center, Seattle, WA 98109; Division of Basic Sciences, Fred Hutchinson Cancer Research Center, Seattle, WA 98109

**Keywords:** Prostate cancer, single cell RNAseq, PTEN, mRNA translation, epithelial cells, immune microenvironment

## Abstract

Advanced prostate malignancies are a leading cause of cancer-related deaths in men, in large part due to our incomplete understanding of cellular drivers of disease progression. We investigated prostate cancer cell dynamics at single-cell resolution from disease onset to the development of androgen independence *in vivo*. We observe a dramatic expansion of a castration-resistant intermediate luminal cell type that correlates with treatment resistance and poor prognosis in human patients. Moreover, transformed epithelial cells and associated fibroblasts create a microenvironment conducive to pro-tumorigenic immune infiltration, which is in part androgen responsive. Androgen independent prostate cancer leads to significant diversification of intermediate luminal cell populations characterized by a range of androgen signaling activity inversely correlated with proliferation and mRNA translation. Accordingly, distinct epithelial populations are exquisitely sensitive to translation inhibition which leads to epithelial cell death, loss of pro-tumorigenic signaling, and decreased tumor heterogeneity. Our findings reveal a complex tumor environment largely dominated by castration-resistant luminal cells and immunosuppressive infiltrates.

## Introduction

Prostate cancer is the most diagnosed malignancy and second leading cause of cancer-related death in men in the United States (Siegel et al., 2021). This is largely due to the development of treatment resistant diseases called castration resistant prostate cancer (CRPC). Historically, CRPC has been considered a singular disease entity. However, this viewpoint has significantly evolved with the advent of next generation sequencing and the characterization of many distinct etiologies (Watson et al., 2015; Bluemn et al., 2017; Labrecque et al., 2019). Intratumoral heterogeneity is also common in prostate cancer, as several studies have highlighted both the complex cellular architecture of the prostate and multiple potential cell-of-origin models (Goldstein et al., 2010; Shen and Abate-Shen, 2010). For instance, while prostate epithelial cells canonically present as either basal cells or highly differentiated luminal cells, a rare but distinct luminal population that primarily resides in the proximal prostate has previously been described and implicated in cancer initiation (Xin et al., 2005; Wang et al., 2006; Korsten et al., 2009; Kwon et al., 2016; Sala et al., 2017; McAuley et al., 2019). This novel cell type was further verified using lineage tracing and single-cell technologies (Liu et al., 2016; Crowley et al., 2020; Joseph et al., 2020; Karthaus et al., 2020; Kwon et al., 2020; Mevel et al., 2020). While the nomenclature for this cell type has been inconsistent, most studies have observed a similar set of biomarkers and characteristics, including increased stemlike and inflammatory/immunogenic signatures (Liu et al., 2016). These luminal cells also appear to fill an important regenerative niche in the prostate environment, including in the context of castration and/or androgen deprivation (Karthaus et al., 2020; Mevel et al., 2020; Kwon et al., 2021). While this suggests they may have a role in castration resistance in the context of cancer, this has yet to be established in CRPC models.

While tumor heterogeneity contributes to treatment resistance, another source of poor clinical outcomes can be found in the consistent lack of response of CRPCs to immunotherapy. Prostate cancer has generally been described as ‘immune-cold’ due to the presence of multiple immunosuppressive cell types (Stultz et al., 2021). For example, tumor infiltration by myeloid-derived suppressor cells (MDSCs) has been implicated as an immunosuppressive phenotype (Garcia et al., 2014; Lopez-Bujanda et al., 2021). Metastatic CRPC also responds poorly to immune checkpoint inhibition and other immunotherapies (Cham et al., 2020). This has been confirmed clinically with autologous active cellular immunotherapy which only demonstrated a small (∼4 month) improvement in survival in patients (Kantoff et al., 2010). Therefore, characterizing the immune microenvironment in advanced prostate cancer may be crucial to better understand this disease and inform potential therapeutic vulnerabilities and/or combinatorial strategies.

Here, we have created an atlas of prostate cellular composition and phenotypic evolution through tumor initiation, progression, and hormone independence using a PTEN loss murine model. We observe a dramatic expansion of the intermediate luminal cell type in cancer and link this epithelial population to treatment resistance in human cohorts. We also characterize increased pro-tumorigenic immune cell recruitment and define cell-cell signaling patterns that contribute to lymphoid and myeloid cell expansion. In addition, using a tissue-specific transgenic model, we demonstrate that cell type specific protein synthesis is essential for the maintenance of tumor heterogeneity in both basal and luminal-intermediate cells in the context of AR-low prostate cancer. Finally, we make our data available for further study through an interactive portal, with the aim of providing a broad resource for the cancer research community. Together, our findings highlight the cell type-specific and patient relevant features of prostate cancer progression and demonstrate the utility of single-cell technologies to uncover novel cell-specific paradigms of treatment resistance.

## Results

### PTEN loss generates differentially-proliferating populations and a transition towards an intermediate luminal cell state in basal cells

To determine the cellular architecture of prostate cancer initiation, we conducted single cell RNA sequencing (scRNAseq) of the ventral prostates of wild-type (WT) and *PB-Cre4;Pten*^*fl/fl*^*;ROSA26-rtTA-IRES-eGFP* (herein referred to as *Pten*^*fl/fl*^) prostate cancer mice (Fig. 1A). In the *Pten*^*fl/fl*^ model, exon 5 of the tumor suppressor PTEN is deleted within basal and luminal epithelial cells of the prostate and mice uniformly develop prostate cancer (Wang et al. Cancer Cell 2003). PTEN is a negative regulator of the oncogenic PI3K-AKT-mTOR signaling pathway which is deregulated in nearly all advance prostate cancer patients (Taylor et al., 2010).

**Fig. 1:**
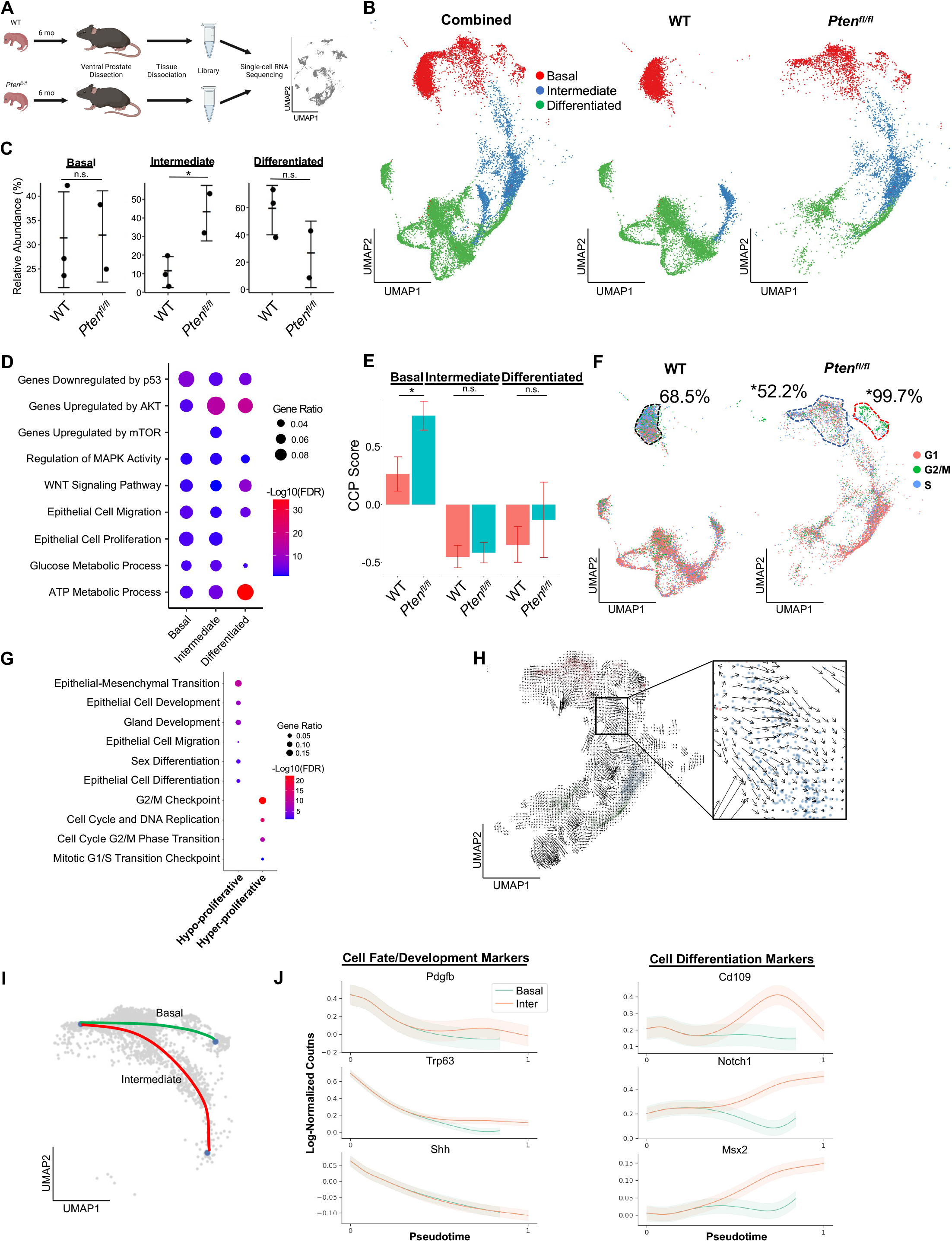
Proliferative Split in Basal Cancer Cells Enables Expansion of Progenitor Cells. **A)** Simplified schematic of single-cell RNA sequencing of WT and *Pten*^*fl/fl*^ ventral prostates. **B)** UMAP of WT and *Pten*^*fl/fl*^ epithelial cells. Left, both conditions superposed; middle, WT only; right, *Pten*^*fl/fl*^ only. Epithelial cell types are demarcated by color (red = basal, green = intermediate, blue = differentiated). **C)** Relative abundance of epithelial cells in WT and *Pten*^*fl/fl*^ mice. Y-axis shows the % composition of each sample by cell type (*p<0.05, negative binomial test). Data presented at +/- SD. **D)** Top GSEA results enriched in *Pten*^*fl/fl*^ compared to WT for each epithelial subtype. All pathways are enriched with FDR < 0.05. **E)** Proliferation signature (CCP) composite score in epithelial cells, clustered by condition (Data presented at +/- SD, *p<0.05, permutation test). **F)** UMAP visualization of cell cycle phase assignment per cell, showing % cells in non-G1 (S or G2/M) (black border = WT basal cells, blue border = hypo-proliferative basal cells in *Pten*^*fl/fl*^, and red border = hyper-proliferative basal cells in *Pten*^*fl/fl*^. *p<0.05, chi-square test). **G)** GSEA between hyper- and hypo-proliferative basal clusters in *Pten*^*fl/fl*^. All pathways are enriched with FDR < 0.05. **H)** RNA velocity analysis of *Pten*^*fl/fl*^ epithelial cells; highlighted section shows intersection of basal and progenitor cells. **I)** Pseudotime trajectories drawn by Palantir through the basal and intermediate compartments, with hypo-proliferating basal cells as the designated start point. **J)** Expression of important cell fate and differentiation regulators along basal-progenitor trajectory. n.s. = not significant.

Quality control and filtering (read count thresholds and excluding cells with high mitochondrial content) resulted in transcriptomes from 24,079 total cells (Table S1A-C). Using SingleR we identified epithelial, stromal, and immune cell types (Aran et al., 2019) (Fig. S1A). To confirm that the epithelial cells in the *Pten*^*fl/fl*^ model underwent Cre-mediated recombination, we measured the frequency of *rtTA-eGFP* fusion mRNA in WT and *Pten*^*fl/fl*^ mice. The *rtTA-eGFP* transcript was present in 14.5% of *Pten*^*fl/fl*^ epithelial cells (Fig. S1B). Given that on average ∼1200 genes/cell were detected and ∼24,000 genes were identified overall, we expect the average transcript to be found in ∼5% of cells in the dataset. Therefore, our finding suggests widespread recombination of PTEN within the epithelial compartment. Importantly, recombination was not observed in WT mice or non-epithelial cell types (Fig. S1B). We further defined the epithelial population via signatures and biomarkers obtained from the Strand (Basal, Urethral, and VP), Sawyers (Basal, L1, and L2), Goldstein (CD38-low), and Xin (Sca-1+) groups (Joseph et al., 2020; Karthaus et al., 2020; Kwon et al., 2016; Liu et al., 2016). Three epithelial cell types were identified in WT and *Pten*^*fl/fl*^ mice: basal *(Krt5+/Sox4+)*, intermediate *(Ppp1r1b+/Clu+/Tactsd2+/Krt4+)*, and differentiated luminal cells (*Nkx3-1+/Sbp+*) (Fig. S1C).

Next, we sought to determine how the epithelial compartment is remodeled in the context of tumor initiation. We observed a dramatic expansion of intermediate luminal cells, as well as a notable decrease in separation between basal and intermediate cells in *Pten*^*fl/fl*^ mice compared to their WT counterparts (Fig. 1B-C). We identified differentially expressed genes (DEG) and performed gene set enrichment analysis (GSEA) on each epithelial subtype, comparing the WT and *Pten*^*fl/fl*^ epithelial compartments (Table S1D). We noted enrichment of oncogene and tumor suppressor pathways in the *Pten*^*fl/fl*^ mice, including upregulation of AKT and mTOR which are expected in this model. We also observed increased *Wnt* signaling, cell migration, and metabolic processes across all 3 epithelial cell types (Fig. 1D). Interestingly, epithelial cell proliferation was enriched in basal and intermediate cells, but not differentiated cells. Given this finding, we hypothesized that the increase in intermediate cell abundance could be caused by higher cell proliferation.

We used the Cell Cycle Progression (CCP) score, a proliferation gene signature developed and validated in human prostate cancer patients (Cuzick et al., 2011), and generated a composite score (see Methods) in our dataset. Surprisingly, while there was a significant increase in CCP score in basal cells in *Pten*^*fl/fl*^ mice compared to WT, we observed no such change in intermediate cells (Fig. 1E). To further characterize epithelial proliferation, we performed cell cycle scoring, assigning one of three phases (G1, G2/M, or S) to each cell in the dataset, and found a striking split in the basal compartment of *PTEN*^*fl/fl*^ mice, but not WT mice (Table S1E). In *Pten*^*fl/fl*^ mouse prostates, 18.6% of basal cells were hyper-proliferative, with 99.7% of these cells in a proliferative phase (G2/M or S phases). The rest of the basal compartment only had 52.2% of cells in a G2/M or S phase, lower than WT basal cells (Fig. 1F, Fig. S1D). We also verified that the increase in basal cell CCP score was specifically due to this hyper-proliferative cluster (Fig. S1E). Another possibility is that a higher recombination efficiency in some clusters led to differential proliferation phenotypes. We analyzed transgene abundance in our basal subclusters and found that in the *Pten*^*fl/fl*^ mouse, 13.6% (12.0%-15.7%) of hypo-proliferating and 19.6% (19.1%-20.3%) of hyper-proliferating basal cells express the *rtTA-eGFP* transgene. This <1.5-fold difference does not account for the >2-fold increase in cycling cells, or the ∼3-fold increase in CCP score observed between the subclusters. As such, we conclude that PTEN loss in the murine prostate promotes a dual phenotype in basal cells, with most cells displaying decreased proliferation while a subset becomes hyper-proliferative.

We further characterized the basal subclusters in *Pten*^*fl/fl*^ mice by performing DEG analysis followed by GSEA (Table S1F). As expected, we observed several cell cycle-related pathways enriched in the hyper-proliferative cluster. However, the hypo-proliferative basal cluster was enriched for several migration, development, and differentiation signatures (Fig. 1G). Transdifferentiation of basal cells to luminal cells in the context of malignant transformation has previously been reported in prostate cancer (Goldstein et al., 2010; Choi et al., 2012). As such, we hypothesized that the hypo-proliferative basal subcluster might transition into intermediate luminal cells, thus providing a potential mechanism for the expansion of this cell type in the absence of increased proliferation. To verify this, we generated pseudotime trajectories using Monocle3 (Cao et al., 2019), which confirmed a direct path from hypo-proliferative basal cells to intermediate luminal cells (Fig S1F). We also performed RNA velocity, a technique to visualize differentiation dynamics on a per cell basis (La Manno et al., 2018). This analysis revealed significant movement from hypo-proliferative, but not hyperproliferative basal cells to intermediate cells (Fig. 1H). Finally, we used a pseudotime algorithm (Setty et al., 2019) to identify two potential trajectories starting from the hypo-proliferative basal cells and ending either in the hyper-proliferative basal cluster or in the intermediate luminal cell compartment (Fig. 1I). We generated clusters of genes with similar expression patterns across these pseudotime axes (Fig. S1G) and performed GSEA on the clusters. We found that several genes involved in cell fate decisions including *Pdgfb, Trp63*, and *Shh* are highly expressed early but rapidly decrease over the trajectories. Conversely, genes implicated in differentiation such as *Notch1* increased over the basal-intermediate path, but not the basal-basal path (Fig. 1J). This suggests that hypo-proliferating basal cells strongly express development markers associated with cell fate choice, but these genes are not expressed in hyper-proliferative basal cells or intermediate cells. Moreover, intermediate cells express higher levels of differentiation-associated genes than either basal subcluster. Together, these findings support the idea that hypo-proliferative basal cells transdifferentiate into intermediate luminal cells during tumorigenesis, while the basal compartment may be replenished by a hyper-proliferative, non-differentiating population.

### *Immune infiltration increases in* Pten^fl/fl^ *mice and is pro-tumorigenic*

Our GSEA of all three epithelial cell types showed an enrichment for immune-related pathways in *Pten*^*fl/fl*^ mice, suggesting increased immunogenic signaling relative to WT mice (Fig. 2A). To determine the impact of epithelial PTEN loss on immune cells, we calculated the relative abundance of immune cells in WT and *Pten*^*fl/fl*^ mice. We found that T cells, macrophages, neutrophils, and dendritic cells were all significantly expanded in the *Pten*^*fl/fl*^ mouse; in fact, only B cells were not expanded compared to WT (Fig. S2A-B). Immune infiltration has previously been characterized as immunosuppressive in PTEN null prostate tumors (Garcia et al., 2014). Therefore, we expected these expanding immune populations to be immunosuppressive and therefore pro-tumorigenic. To test this hypothesis, we assigned activation states or subtypes to immune cells based on known biomarkers (Fig. 2B, Fig. S2C-E). First, we characterized the neutrophil population as myeloid-derived suppressor cells (MDSCs) based on published biomarkers (Alshetaiwi et al., 2020) (Fig. S2C). We also found three macrophage cell states: tumor-associated macrophages (TAMs), M2-activated macrophages, and M1-activated macrophages. Interestingly, the M1 cells expressed the AR-dependent markers *Sbp* and *Defb50* (Fig. S2D). As a result, we termed them tissue-resident macrophages (TRM). Finally, we characterized three T cell subtypes: CD8^+^ T cells, gamma-delta T cells, and natural killer T (NKT) cells. CD8^+^ and gamma-delta T cells expressed the markers of exhaustion and immunosuppression *Pdcd1* and *Ctla4*, suggesting their cytotoxic activity may be dampened in the context of PTEN loss (Fig. S2E). All of these cell types except for NKT cells and TRMs are expanded in *Pten*^*fl/fl*^ mice, implying a potential role for these immune cells in establishing or maintaining a pro-tumorigenic prostate tumor microenvironment (Fig. 2C).

**Fig. 2:**
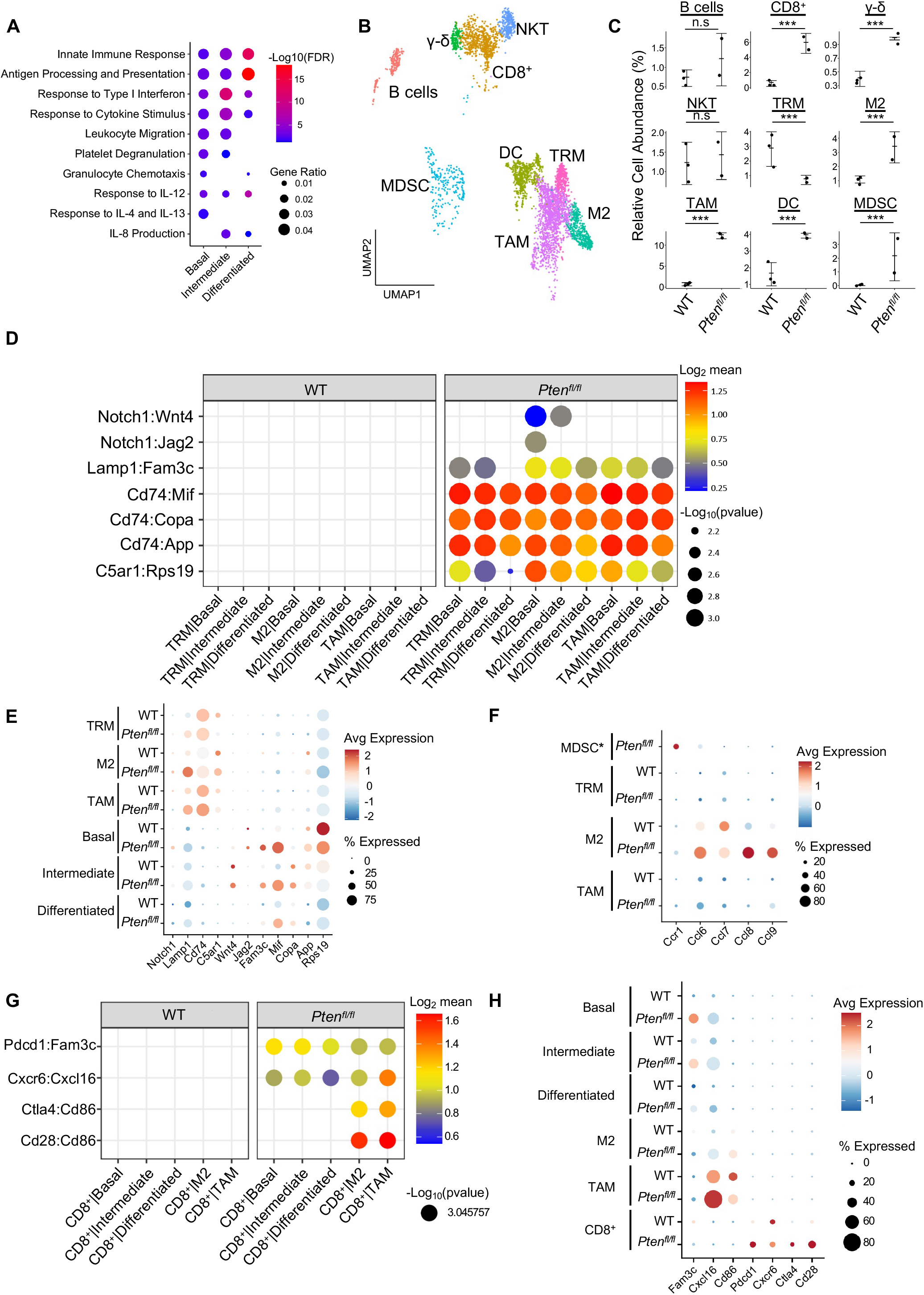
Immune Recruitment in *Pten*^*fl/fl*^ Prostates is Mediated by both Epithelial and Immune Cell Signaling. **A)** Top immune-related GSEA results enriched in *Pten*^*fl/fl*^ compared to WT mice each epithelial subtype. All pathways are enriched with FDR < 0.05. **B)** UMAP visualization of immune cells labeled by cell subtype or state. **C)** Relative abundance of immune cells in WT and *Pten*^*fl/fl*^ mice. Y-axis shows the % composition of each sample by cell type (Data presented at +/- SD, ***p<0.001, negative binomial test). **D)** Dot plot of signaling interactions between macrophages and epithelial cells. Y-axis, ligand-receptor pairs from CellphoneDB database. X-axis, cell-cell pairings. Interactions are directional: the first gene in a pair is expressed in the first cell in the cell-cell interaction. **E)** Dot plot of epithelial ligand and macrophage receptor gene expression in WT and *Pten*^*fl/fl*^ mice. **F)** Dot plot of Ccr1 and Ccr1 ligand expression in MDSCs and macrophages in WT and *Pten*^*fl/fl*^ ventral prostates. MDSCs are only present in *Pten*^*fl/fl*^ and therefore do not have a WT row (denoted by asterisk). **G)** Plot of signaling interactions between CD8 T cells and epithelial cells and macrophages. **H)** Dot plot of CD8^+^ T cell receptors and epithelial and macrophage ligand gene expression in WT and *Pten*^*fl/fl*^ ventral prostates. N.s. = not significant.

M2 macrophages, TAMs, and MDSCs are generally pro-tumorigenic (Zaynagetdinov et al., 2011; Chanmee et al., 2014; Garcia et al., 2014). Together with exhausted CD8^+^ T cells, this suggests a broadly pro-tumorigenic immune environment. We hypothesized that cell-cell signaling originating from tumor epithelial cells may play an important role in recruiting immune cells to the prostate. To probe cell-cell interactions in our system, we used a ligand-receptor database and interaction algorithm that classifies the strength of specific ligand-receptor interactions between cell groups (Efremova et a., 2020). We found key interactions between epithelial subtypes and M2 macrophages and TAMs, such as interactions targeting the CD74 receptor, that point to active recruitment (Fig. 2D, Table S2). Epithelial ligands were more highly expressed in *Pten*^*fl/fl*^ mice than in WT mice, which corresponds to the increased macrophage abundance in *Pten*^*fl/fl*^ mice (Fig. 2E).

Additionally, we observed increased CCL2/7/11-CCR2 interactions between fibroblasts and M2/TAMs upon PTEN loss, suggesting that fibroblasts in prostate cancer may also play an active role in macrophage recruitment (Fig. S2F). CCL2/7/11 are all significantly upregulated in *Pten*^*fl/fl*^ fibroblasts (Fig. S2G). Interestingly, CCR2 is expressed highly in M2 and TAMs in cancer, but not in TRMs (Fig. S2H). The lack of fibroblast signaling to TRMs provides a possible cellular explanation for the lower abundance of this macrophage subtype in *Pten*^*fl/fl*^ mice compared to WT. Another possibility is that TRMs polarize and transition into M2/TAM macrophages, potentially in response to epithelial signaling. Overall, these findings highlight a role for epithelial cells and fibroblasts in recruiting pro-tumorigenic macrophages to the prostate in the context of PTEN loss.

While we observed significant epithelial to macrophage signaling, the top interactions between epithelial cells and MDSCs were characterized by receptors expressed in epithelial cells and ligands expressed in MDSCs. As such, it is unlikely that epithelial cells are responsible for the expansion of MDSCs in the prostate (Fig. S2I). Instead, we noted increased expression of the CCR1 receptor in MDSCs, and a concomitant increase in CCL6/7/8/9 expression in M2 macrophages. These genes are known CCR1 ligands (Korbecki et al., 2020), which suggests that MDSCs are recruited by M2-activated macrophages (Fig. 2F). Interestingly, CCL6/7/8/9 are not expressed in TRMs or TAMs, highlighting the specificity of M2-MDSC signaling.

Finally, we examined interactions between epithelial cells and T cells. We found both attractive (*Cxcr6*-*Cxcl16*) and pro-exhaustion (*Pdcd1*-*Fam3c*) interactions between CD8^+^ T cells and epithelial cells. We found these same interactions between macrophages and CD8^+^ cells, as well as *Cd86*-*Ctla4*/*Cd28* interactions. CTLA4 and CD28 belong to the same family of T cell co-stimulator receptors. CTLA4 has a suppressive effect while CD28 is activating, and both receptors are targeted by the CD86 ligand (Gmunder et al., 1984; Salomon and Bluestone, 2001; Boesteanu and Katsikis, 2009). These findings imply a synergistic signaling pattern with both tumor epithelia and macrophages recruiting and mitigating the cytotoxicity of T cells (Fig. 2G). We noted that TAMs exhibited particularly robust expression of Cxcl16 and Cd86, suggesting that similarly to M2-mediated MDSC recruitment, TAMs may specifically be responsible for modulating T-cell abundance and function (Fig. 2H).

Overall, our data suggests that pro-tumorigenic macrophages are recruited early on in cancer by epithelial and fibroblast signaling. These macrophages then assist tumor signaling in remodeling the immune environment, including recruiting and exhausting cytotoxic CD8^+^ T cells and attracting pro-tumorigenic MDSCs. Given ligand expression data, M2 macrophages may be mainly responsible for MDSC recruitment and TAMs for CD8^+^ T cell recruitment. These findings reveal a complex signaling system with multiple coordinated sources of ligand expression working in tandem to build a microenvironment favorable to tumor escape from immunological suppression.

### Castration-induced intermediate cell heterogeneity drives resistance to androgen deprivation

Having examined epithelial and immune populations in prostate cancer initiation, we sought to determine how these cells reorganize over the course of castration resistance. It has been shown that castration of *Pten*^*fl/fl*^ mice leads to the emergence of AR-low and proliferative tumors (Liu et al., 2019). As such, we conducted scRNAseq in *Pten*^*fl/fl*^ mice with and without castration and evaluated the changes that occurred in the epithelial compartment (Fig. 3A). Castration caused the intermediate luminal cell population to expand while the differentiated luminal cells disappeared entirely (Fig. 3B-C). We hypothesized that androgen deprivation may differentially affect epithelial subtypes. To investigate this, we generated an AR activity score using a 20-gene signature (Hieronymus et al., 2006) and found that intermediate cells had high AR activity in WT prostate, but very low activity in both intact and castrated PTEN-null conditions. On the contrary, differentiated cells retained high AR activity in *Pten*^*fl/fl*^ mice (Fig. 3D). This suggests that loss of PTEN in luminal cells decreases their reliance on AR signaling and induces expansion of the intermediate cells that can survive androgen deprivation. Notably, there were no significant changes in proliferation between the intact and castrate conditions in basal and intermediate cells (Fig. S3A).

**Fig. 3:**
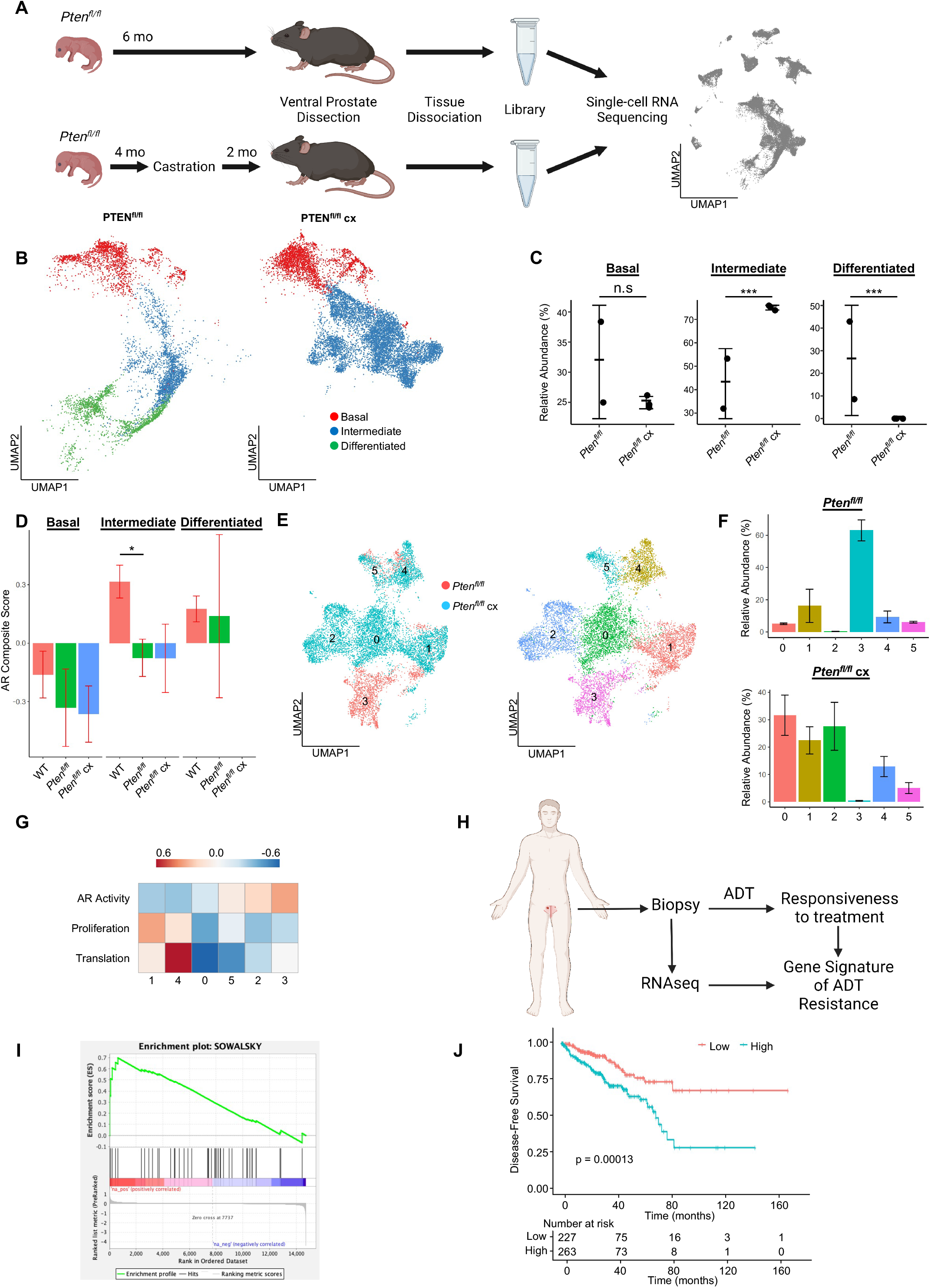
Progenitor Cells are Primed for Survival in the Context of Castration and Correlate to Treatment Resistance. **A)** Simplified schematic of setup for single-cell sequencing of *Pten*^*fl/fl*^ intact and *Pten*^*fl/fl*^ castrated (cx) ventral prostates. **B)** Split UMAP visuzalizations of *Pten*^*fl/fl*^ and *Pten*^*fl/fl*^ cx epithelial cells. **C)** Relative abundance of epithelial cells in *Pten*^*fl/fl*^ intact and cx prostates. Y-axis shows the % composition of each sample by cell type (Data presented at +/- SD, ***p<0.001, negative binomial test). **D)** Androgen Receptor (AR) gene signature composite score in epithelial cells, clustered by condition (Data presented at +/- SD, *p<0.05, permutation test). **E)** UMAP visualization of intermediate cells in *Pten*^*fl/fl*^ intact and cx prostates. Left, colored by condition; right, colored by clusters 0-5. **F)** Relative abundance of intermediate clusters. Top, intact *Pten*^*fl/fl*^; bottom, *Pten*^*fl/fl*^ cx (Data presented at +/- SD). **G)** Heatmap of composite score for AR, CCP, and Reactome translation gene signatures in intermediate clusters. **H)** Diagram of clinical trial used to establish gene signature of androgen deprivation treatment resistance (NCT02430480). **I)** Enrichment plot of ADT resistance gene signature relative to progenitor cell DEGs between *Pten*^*fl/fl*^ and *Pten*^*fl/fl*^ cx (adjusted p-value = 0.00381). **J)** Kaplan-Meier curve of disease-free survival for prostate cancer patients in TCGA database with or without high RNA expression of top correlated genes from **Fig. 3H**. Red line, patients with normal expression of all genes; blue line, patients with expression of at least 1 gene with TPM in the 80^th^ percentile or above. n.s. = not significant.

Given the increase in intermediate cells associated with castration resistant tumor growth, we next asked how castration modulates phenotypic diversity in this population. We isolated and re-clustered intermediate cells from *Pten*^*fl/fl*^ intact and castrated mice and found six distinct clusters (Fig. 3E). We found that the majority of intact intermediate cells (62.8%) congregated in a single cluster (cluster 3), while castrated cells were widely distributed over four unique groups (clusters 0,1,2, and 4) (Fig. 3F). DEG analysis showed high expression of AR-dependent genes *Sbp, Defb50*, and *Spink1* in cluster 3, suggesting this cell population may have high AR signaling activity relative to other intermediate cells (Fig. S3B, Table S3A-F). In addition, we observed high expression of the basal cell markers *Krt5* and *Krt15* in cluster 1; this cluster corresponds to the intermediate cells proximal to basal cells on the epithelial UMAP. This supports our hypothesis that some intermediate cells are derived from basal transdifferentiation in the context of cancer. We also noted multiple ribosomal genes upregulated in clusters 1 and 4, suggesting increased translation in these clusters (Fig. S3B). GSEA confirmed enrichment of multiple translation pathways in clusters 1 and 4 (Fig. S3C). We also investigated whether castration leads to increased translation rates in basal cells. We found that several genes encoding ribosomal proteins were overexpressed in castrated basal cells compared to intact *Pten*^*fl/fl*^ mice (Fig. S3D). Together, these findings demonstrate that decreasing AR signaling promotes increased heterogeneity of the intermediate cell populations and potentially diversifies cell type specific translation dependence in both basal and intermediate cells in the context of PTEN loss.

To further characterize the functional differences in castration resistant intermediate cells, we generated AR activity, proliferation, and translation scores for each cluster. Although AR activity is greatly decreased in these cells in cancer, we found a gradient of residual AR signaling activity, with cluster 3 having the highest score (Fig. 3G). Importantly, proliferation and translation activity scores were inversely correlated with AR signaling, with clusters 1 and 4 exhibiting high proliferation and translation scores and low AR scores (Fig. 3G). We designated the clusters as AR-high, -medium, or -low and noted that while intact *Pten*^*fl/fl*^ mouse prostates contain mostly AR-high intermediate cells, ∼25% of the compartment is AR-low (Fig. S3E-F). These findings suggest that castration selected for existing AR-low, highly proliferative regions of the intermediate luminal cells, conserving some “intact-like” regions with relatively high AR activity and low proliferation.

Given the widespread use of the *Pten*^*fl/fl*^ model (Ding et al., 2011; Svensson et al., 2011; Hsieh et al., 2012; Garcia et al., 2014; Hsieh et al., 2015; Ku et al., 2017; Allott et al., 2018; Antoch et al., 2020; Morel et al., 2021; Quaglia et al., 2021) and our new understanding of cellular dynamics in the context of disease progression, we sought to determine which cell type and context most closely correlated with human prostate cancers that went on to resist androgen deprivation therapy (ADT). To this end we used a gene signature of ADT resistance derived from prostate tumors prior to treatment with ADT plus the AR inhibitor enzalutamide, in a neoadjuvant setting (Fig. 3H) (NCT02430480) (Karzai et al., 2020; Wilkinson et al., 2021; Ku et al., 2021). We generated DEG lists for each epithelial cell type comparing WT and *Pten*^*fl/fl*^ or *Pten*^*fl/fl*^ intact and castrated mice and performed enrichment analysis using the ADT resistance signature. Out of all the cell types, castrated intermediate cells compared to intact intermediate cells exhibited the most enrichment for the resistance signature (Fig. 3I). The top five genes from the resistance signature that were present in castrated intermediate cells were ATP1B1, BST2, CP, IGFBP3, and PTTG1. Importantly, these genes were downregulated in cluster 3 of our intact intermediate cells compared to castrate clusters demonstrating specificity for aggressive disease (Fig. S3G). Furthermore, all 5 genes were upregulated in human tumors that exhibited pathologic resistance to ADT (Fig. S3H). Lastly, we sought to investigate whether these genes were associated with worse outcomes for prostate cancer patients. We examined disease-free survival (DFS) of patients stratified by high or low expression of the five top genes across two major prostate cancer cohorts (The Cancer Genome Atlas Research Network, 2015; Taylor et al., 2010). We found that patients whose tumor samples expressed high levels of any of these five most enriched genes experienced significantly shorter DFS (Fig. 3J, S3I). Together, our findings demonstrate that prostate cancer progression in the context of PTEN loss and castration resistance is associated with the expansion of intermediate cell diversity which closely correlates with worse outcomes and treatment resistance in prostate cancer patients.

### Androgen deprivation decreases immune cell abundance but activates TNF signaling

Androgen deprivation induces a host of physiological changes in the prostate, including modulations of immune signaling (Sha et al., 2015; Lopez-Bujanda et al., 2021). Having observed significant epithelial changes in castrated *Pten*^*fl/fl*^ mice, we next investigated the consequences of castration on the immune environment. We observed a significant decline in the abundance of all 3 macrophage subtypes, as well as CD8^+^ T cells, relative to intact *Pten*^*fl/fl*^ mice (Fig. 4A-B). We performed ligand-receptor interaction analysis to understand how androgen deprivation disrupts cell-cell signaling patterns (Table S4). We found that epithelial signaling to macrophage cells was still intact, with relatively little change in ligand or receptor expression (Fig. S4A-B). However, the CCL2/7/11-CCR2 signaling axis from fibroblasts to M2 macrophages and TAMs was entirely ablated in castrated *Pten*^*fl/fl*^ mice (Fig. 4C). CCR2 expression was significantly decreased in macrophages, and Ccl2/7/11 were all dramatically decreased in fibroblasts (Fig. 4D). This suggests that fibroblast-mediated macrophage recruitment is interrupted by androgen deprivation. Indeed, androgen signaling is known to promote pro-tumorigenic macrophage function as well as macrophage recruitment via CCR2 expression, lending credence to this hypothesis (Lai et al., 2009; Cioni et al., 2020; Becerra-Diaz et al., 2020). TRMs also decreased in abundance, but they were not targeted by fibroblast signaling and epithelial signaling was uninterrupted. Given the expression of AR-dependent genes in this macrophage subtype (Fig. S2D), we speculated that loss of androgen signaling could be deleterious to this population. Indeed, in intact *Pten* ^*fl/fl*^ mice TRMs had high AR activity relative to other macrophage subtypes, and this activity was decreased by castration (Fig. S4C). This suggests that castration interrupts fibroblast-mediated pro-tumorigenic M2 and TAM recruitment and depletes the androgen-dependent tissue-resident macrophage reservoir, leading to decreased macrophage abundance in the prostate.

**Fig. 4:**
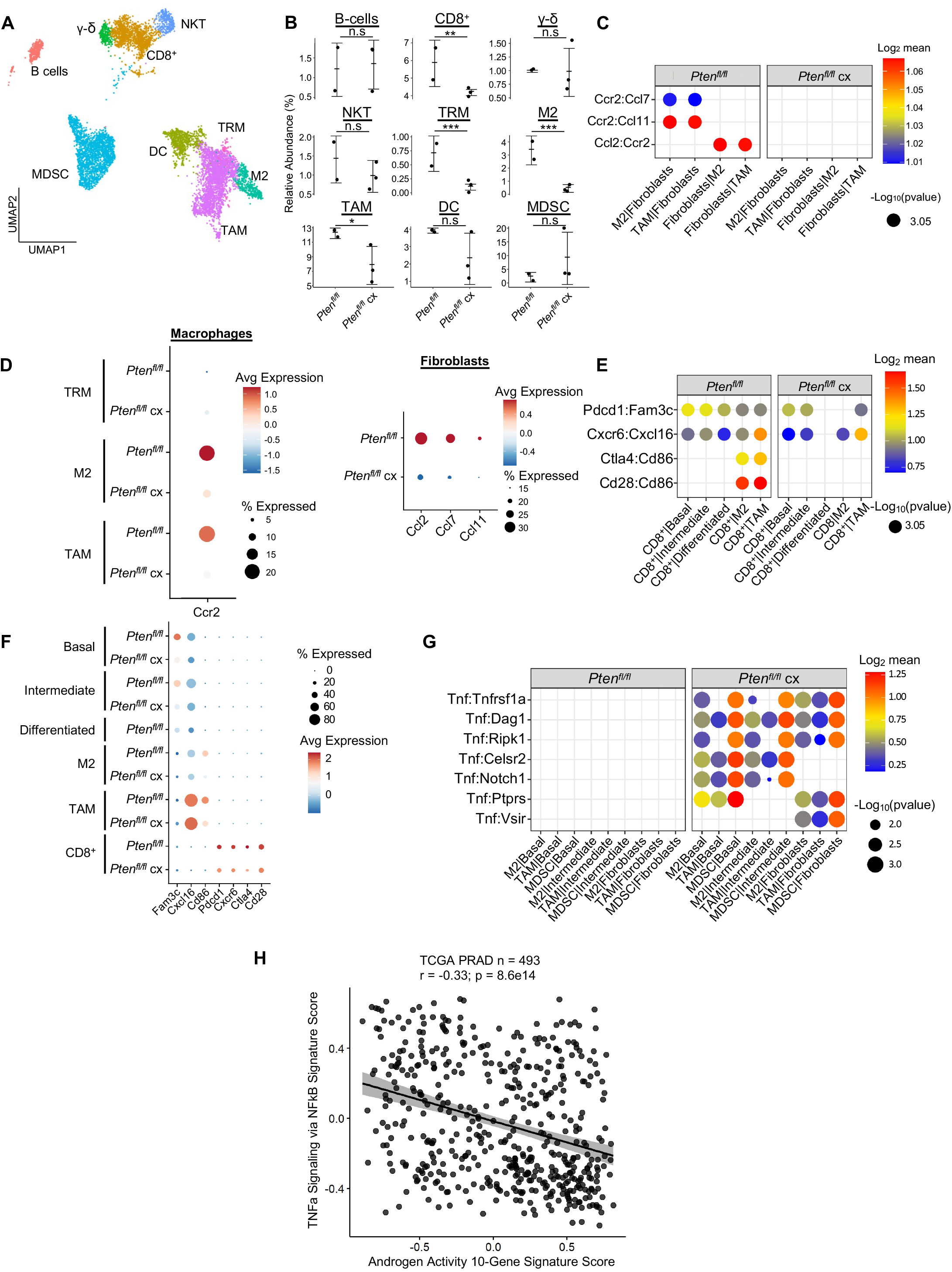
Castration Remodels Immune Environment via Fibroblast Signaling and Increases TNF Pathway Activity. **A)** Combined UMAP visualization of immune cells in *Pten*^*fl/fl*^ and *Pten*^*fl/fl*^ cx ventral prostates. **B)** Relative abundance of immune cells in *Pten*^*fl/fl*^ intact and cx mice. Y-axis shows the % composition of each sample by cell type (Data presented at +/- SD, *p<0.05, **p<0.01, ***p<0.001, negative binomial test). **C)** Dot plot of signaling interactions between macrophages and fibroblasts. **D)** Dot plot of *Ccr2* expression in M2 macrophages and TAMs (left). Dot plot of *Ccr2* ligand expression in fibroblasts in *Pten*^*fl/fl*^ intact and cx mice (right). **E)** Dot plot of signaling interactions between CD8^+^ T cells and epithelial and macrophage cells in *Pten*^*fl/fl*^ intact and cx mice. **F)** Dot plot of epithelial and macrophage ligands and CD8^+^ T cell receptor gene expression in *Pten*^*fl/fl*^ intact and cx mice. **G)** Dot plot of TNF signaling interactions between myeloid and epithelial/fibroblast cells in *Pten*^*fl/fl*^ intact and cx prostates. **H)** Scatter plot of TCGA PRAD study patient signature composite scores. Y-axis, TNF signaling signature score; X-axis, AR signaling signature score (Pearson’s correlation).

Macrophages likely contribute to CD8^+^ T cell recruitment in intact *Pten*^*fl/fl*^ mice (Fig. 2G-H); we speculated that macrophage-mediated signaling might be interrupted in the context of castration and cause a decrease in CD8^+^ T cell abundance. Indeed, while epithelial-CD8^+^ interactions were mostly intact, CD86-CD28/CTLA4 signaling from M2 macrophages and TAMs was disrupted, and CD86 expression was greatly reduced in both M2s and TAMs (Fig. 4E). In addition, receptor expression was decreased in CD8 T-cells, including suppressive markers CTLA4 and PDCD1 (Fig. 4F). These findings suggest that depletion of the macrophage population causes a decrease in both CD8^+^ T cell recruitment and suppression, possibly leading to more cytotoxic but less abundant CD8^+^ T cells in castrated *Pten*^*fl/fl*^ mice.

Finally, TNF signaling has previously been implicated as a pro-tumorigenic factor in AR low prostate cancer (Mizokami et al., 2000; Sha et al., 2015). Accordingly, we examined TNF interactions in our ligand-receptor analysis and found a striking enrichment of TNF pathway activity in castrated mice. Specifically, pro-tumorigenic myeloid cells (M2 macrophages, TAMs, and MDSCs) expressing TNF interact with multiple receptors in epithelial cells and fibroblasts (Fig. 4G). To investigate whether this association held true in human prostate cancer, we correlated a 200 gene signature of TNF activity (Griss et al., 2020) with AR signaling activity in the TCGA dataset (The Cancer Genome Atlas Research Network, 2015).We found a significant inverse correlation between TNF and AR activity in human patients, validating our finding that TNF signaling is induced in prostate cancer upon castration (Fig. 4H). This correlation also held true when only considering patients with PTEN loss (Fig. S4D). We conclude that castration in the *Pten*^*fl/fl*^ mouse provokes several large-scale cellular signaling changes that result in decreased macrophage and CD8^+^ T cell populations and increased TNF signaling.

### Translation inhibition in AR-low prostate cancer is lethal to basal and intermediate cells and disrupts pro-tumorigenic signaling

High mRNA translation rates have previously been associated with aggressive AR-low prostate cancer (Liu et al., 2019). However, understanding how the per cell requirement for aberrant translation enables tumor heterogeneity has been technically challenging. Therefore, it remains to be determined which prostate cancer epithelial cell types require increased translation for androgen independent growth. Given the strong correlation between proliferation and translation observed in both basal and intermediate cells (Fig. 3G, Fig. S3F), we hypothesized that inhibiting translation in the *Pten*^*fl/fl*^ mouse could be deleterious to both cell types. To investigate this possibility, we used the *PB-Cre4;Pten*^*fl/fl*^*;ROSA26-rtTA-IRES-eGFP;TetO-4ebp1*^*M*^ mouse model (herein referred to as *Pten*^*fl/fl*^*;4ebp1*^*M*^). In this model, Cre-mediated recombination leads to PTEN loss and expression of the rtTA protein in both basal and intermediate cells (Fig. S1B). When mice are treated with doxycycline, a mutant *4ebp1* allele (*4ebp1*^*M*^) is expressed (Hsieh et al., 2015). 4EBP1 is a negative regulator of translation initiation and functions via inhibition of eIF4F complex assembly (Schuster and Hsieh, 2019). This mutant allele cannot be inactivated via mTOR-mediated phosphorylation and its expression robustly inhibits eIF4F complex formation and translation initiation in prostate epithelia (Fig. 5A). Using this model, we sought to determine the epithelial cell type specific dependencies of castration resistant prostate cancer.

**Fig. 5:**
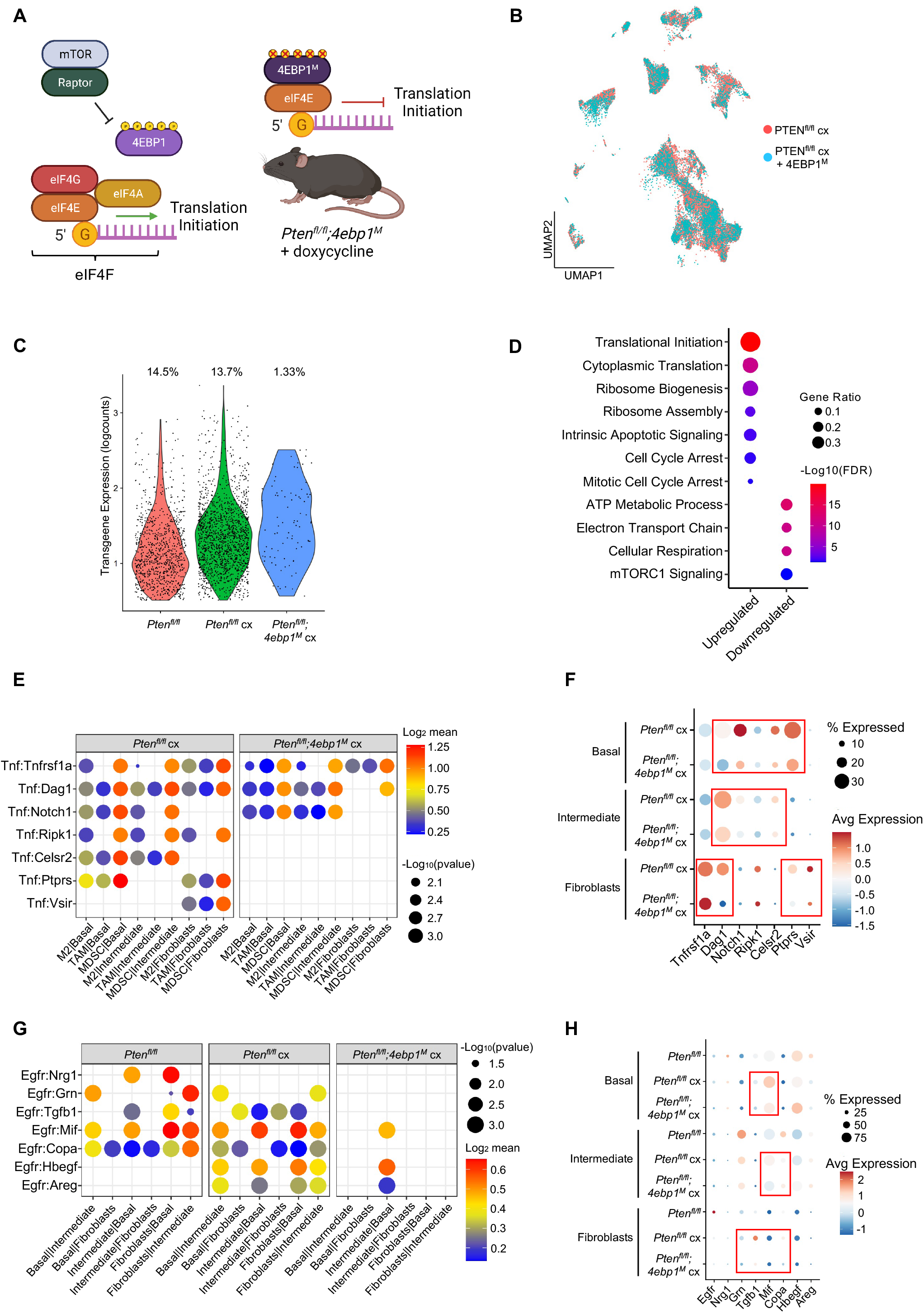
4EBP1^M^ Expression is Lethal in Epithelial Cells and Decreases EGFR and TNF ligands in Epithelial and Fibroblasts. **A)** Simplified schematic of the eIF4F translation initiation complex and how the 4ebp1^M^ protein functions in the *Pten*^*fl/fl*^*;4ebp1*^*M*^ mouse model when treated with doxycycline. **B)** UMAP visualization of epithelial cells in *Pten*^*fl/fl*^ cx and *Pten*^*fl/fl*^*;4ebp1*^*M*^ cx prostates, colored by genotype. **C)** Violin plot of *rtTA-eGFP* transgene expression in epithelial cells in each *Pten*^*fl/fl*^ condition. Plot shows only cells expressing the transgene; each dot represents a cell. Percentages represent the proportion of transgene-positive cells in each condition. **D)** Dot plot of top GSEA results from DEG analysis of transgene-positive basal cells in *Pten*^*fl/fl*^*;4ebp1*^*M*^ cx mice compared to *Pten*^*fl/fl*^ cx ventral prostates. All pathways are enriched with FDR < 0.05. **E)** Dot plot of TNF signaling interactions between myeloid and epithelial/fibroblast cells in *Pten*^*fl/fl*^ cx and *Pten*^*fl/fl*^*;4ebp1*^*M*^ cx mice. **F)** Dot plot of TNF and TNF ligand expression in myeloid cells, epithelial cells, and fibroblasts in *Pten*^*fl/fl*^ cx and *Pten*^*fl/fl*^*;4ebp1*^*M*^ cx prostates. Red boxes highlight ligands with decreased expression in *Pten*^*fl/fl*^*;4ebp1*^*M*^ mice. **G)** Plot of EGFR signaling interactions between epithelial cells and fibroblasts in *Pten*^*fl/fl*^ cx and *Pten*^*fl/fl*^*;4ebp1*^*M*^ cx prostates. **H)** Dot plot of EGFR and EGFR ligand expression in epithelial cells and fibroblasts in *Pten*^*fl/fl*^ cx and *Pten*^*fl/fl*^*;4ebp1*^*M*^ ventral prostates. Red boxes highlight ligands with decreased expression in *Pten*^*fl/fl*^*;4ebp1*^*M*^ mice.

Initially, no major differences were observed in the UMAP comparing all epithelial cells in castrated *Pten* ^*fl/fl*^ mice with or without *4ebp1*^*M*^ (Fig. 5B). However, we noted a striking (∼10-fold) depletion of the *rtTA-eGFP* transgene which is required for 4EBP1^M^ induction in *Pten*^*fl/fl*^*;4ebp1*^*M*^ basal and intermediate epithelial cells compared to the *Pten*^*fl/fl*^ model (Fig. 5C, Table 1). We hypothesized that translation inhibition in AR-low prostate cancer might be lethal or confer a competitive disadvantage to AR low epithelial cells, and that the bulk of the remaining epithelia did not express the *4ebp1*^*M*^ and were simply castrated cells that did not express the transgene. To test this hypothesis, we performed DEG analysis only on transgene-positive cells, comparing castrated cells with or without *4ebp1*^*M*^. We found many more differentially expressed genes between these groups in basal cells (465 differentially expressed genes) than in the non-filtered analysis (56 DEGs) (Table S5A-B). We did not observe a significant change in the number of DEGs between castrated *Pten*^*fl/fl*^ and *Pten*^*fl/fl*^*;4ebp1*^*M*^ intermediate cells when filtering for *rtTA-eGFP*+ cells. This may be due to the high phenotypic diversity in this compartment and the very low proportion of transgene-positive intermediate cells after 4EBP1^M^ induction (<1%, Table 1) causing a lack of robustness in the DEG analysis. However, we did observe a significant increase in the expression of translation regulators such as ribosomal proteins of the small and large subunits and a translation elongation factor (Table S5B), which may represent a stress response to eIF4F inhibition.

**Table 1:** Transgene abundance in PTEN mouse epithelia

| Cell Type |  | *Pten*<sup>fl/fl</sup> | *Pten*<sup>fl/fl</sup> cx | *Pten*<sup>fl/fl</sup>;4ebp1<sup>M</sup> |
| --- | --- | --- | --- | --- |
| Basal | All | 1795 | 2718 | 1565 |
|  | *rtTA-eGFP*+ | 255<br>(14.2%) | 422<br>(15.5%) | 41<br>(2.6%) |
| Intermediate | All | 2391 | 8230 | 4760 |
|  | *rtTA-eGFP*+ | 438<br>(18.3%) | 1073<br>(13.0%) | 43<br>(0.90%) |

Interestingly, upon performing pathway analysis on basal cell DEGs we found enrichment of pathways relating to translation, cell cycle arrest, and apoptosis, and observed downregulation of mitochondrial function and mTORC1 signaling pathways (Fig. 5D). These findings suggest that remaining transgene-positive basal cells in the *Pten*^*fl/fl*^*;4ebp1*^*M*^ mice may be undergoing cell cycle arrest, interruption of growth processes such as the mTOR pathway, and apoptosis, as well as compensating for eIF4F inhibition by increasing transcription of translational machinery components. We also noted that in the basal compartment, the proportion of hyperproliferative cells had decreased drastically (Table 2). Given the small number of transgene-positive cells in these samples, this change may be stochastic rather than biological, but this suggests that highly proliferative basal cells are more dependent upon high translation than less proliferative basal cells.

**Table 2:** Basal proliferative subset proportions in *rtTA-eGFP*+ cells

| Basal Subset | Hypo-proliferative | Hyper-proliferative |
| --- | --- | --- |
| *Pten*<sup>fl/fl</sup> | 188 (73.7%) | 60 (23.5%) |
| *Pten*<sup>fl/fl</sup> cx | 313 (74.1%) | 98 (23.2%) |
| *Pten*<sup>fl/fl</sup>;4ebp1<sup>M</sup> | 38 (97.4%) | 1 (2.6%) |

Given the large-scale changes in epithelial populations caused by 4EBP1^M^ induction in castrated *Pten*^*fl/fl*^ mice, we next asked how cell-cell signaling between the remaining epithelial cells and other compartments were affected by the loss of basal and intermediate cell populations through translation inhibition (Table S5C). We found that TNF signaling was greatly decreased between myeloid and epithelial cells (Fig. 5E-F). In addition, epithelial-fibroblast and inter-epithelial EGFR signaling also decreased significantly (Fig. 5G-H). EGFR activity is associated with worse cancer prognosis, and inhibition of EGFR signaling has been proposed as a therapeutic approach in advanced prostate cancer (Kim et al., 2006; Guerin et al., 2010; Xiong et al., 2020). Overall, these findings show that both basal and intermediate cancer cell types require increased translation initiation to maintain castration resistance and that aberrant mRNA translation is required for tumor heterogeneity. Furthermore, translation inhibition of distinct epithelial populations can impact the local tumor microenvironment.

## Discussion

Intermediate luminal cells are an important niche cell type in prostate epithelia with roles in inflammation, regeneration, and tumor progression (Liu et al., 2016; Kwon et al., 2016; Karthaus et al., 2020). While these cells have been an active area of study in recent years, there remain multiple questions on their role in healthy and diseased tissue. Here, we demonstrated that PTEN deletion in mouse prostate epithelial cells results in expansion of the intermediate luminal subtype, which is partly mediated by the transdifferentiation of a specific basal cell population. Interestingly, this transdifferentiation phenotype was entirely absent in the WT mouse, where the basal and luminal compartments clustered separately. This corresponds well with lineage tracing studies by Choi et al. (2012) who showed that basal cells can transition to luminal phenotypes prior to transformation. Together these findings suggest that intermediate cells play an important role in tumorigenesis, potentially acting as a newly-uncovered transition point for transforming basal cells.

We also validate previous findings that intermediate cells are castration resistant (Mevel et al., 2020; Joseph et al., 2020) and propose a “priming” model in which androgen signaling is drastically decreased in these cells even in intact cancer contexts, permitting survival upon androgen deprivation. We also observe a significant increase in intermediate cell heterogeneity upon castration. This finding suggests that specific portions of the intermediate compartment are responsible for the increased proliferation previously observed in castrated *Pten*^*fl/fl*^ mice (Liu et al., 2019) and that intermediate cells with less AR signaling are more likely to be highly proliferative. We further corroborate that high translation rates may be required to achieve this high-proliferation phenotype. Finally, we note that the AR-low, proliferation-high regions express basal markers, indicating that basal cells that transition to intermediate cells in tumorigenesis may significantly contribute to castration resistance and hyper-proliferation. Given the presence of intermediate cells in WT prostates and this novel basal phenotypic switch, we hypothesize that the intermediate cell compartment is highly heterogeneous in cancer partly due to multiple cellular origins converging on a common phenotype. Existing intermediate cells likely make up a portion of intermediate tumor cells and may be less proliferative than newly differentiated basal cells. In addition, the ablation of differentiated luminal cells upon castration could be due to cell death or to de-differentiation leading to a more intermediate-like cell state. We conclude that intermediate cells in cancer are highly plastic and may originate from multiple sources, resulting in functional heterogeneity with notable consequences for cancer proliferation and castration resistance.

The prostate tumor environment is typically described as immunosuppressive and does not readily respond to immunotherapies (Kantoff et al., 2010; Cham et al., 2020; Dong et al., 2021). Correspondingly, we show that the immune environment of the *Pten*^*fl/fl*^ mouse prostate is highly enriched for pro-tumorigenic immune cells, specifically immunosuppressive myeloid cells and exhausted CD8^+^ T cells. We carefully delineated cell-cell signaling patterns and found that epithelial cells, fibroblasts, and macrophages contribute to immune recruitment. Since M2 and tumor-associated macrophages most significantly contribute to recruitment of other immune cells, macrophages are likely activated and recruited first during tumorigenesis, and subsequently emit chemokines and other ligands that help attract and exhaust CD8^+^ T cells as well as pro-tumorigenic MDSCs. Interestingly, specific macrophage subtypes seem differentially responsible for distinct recruitment patterns: M2 macrophages express very high levels of MDSC-associated chemokines, while TAMs recruit CD8^+^ T cells via high expression of CXCL16 and CD86. Given these results, interrupting the recruitment of tumor-associated macrophages may be a valid strategy for depleting pro-tumorigenic immune populations and overcoming immunotherapy resistance in prostate cancer.

Androgen deprivation in the *Pten*^*fl/fl*^ mouse leads to a decrease in macrophage and CD8^+^ T cell abundance, likely due to ablation of fibroblast-mediated chemokine signaling. Androgen signaling is active in tissue-resident macrophages and can induce pro-tumorigenic behaviors including increased migration and proliferation in prostate cancer cells (Cioni et al., 2020). In addition, AR can promote CCR2 expression and facilitate macrophage activity and recruitment (Lai et al., 2009). This corresponds well to our finding that fibroblast-mediated signaling towards CCR2 contributes to macrophage recruitment and is interrupted upon castration, and suggests that androgen deprivation decreases macrophage abundance in the prostate tumor environment. Conversely, castration increased TNF signaling from myeloid cells to epithelial cells and fibroblasts. TNF signaling has been described as pro-tumorigenic in AR-low prostate cancer (Mizokami et al., 2000; Sha et al., 2015), and we confirmed that TNF activity was inversely associated with AR activity in human patients. This paradigm supports the role of macrophages and neutrophils in maintaining a favorable tumor environment even in the context of androgen loss and suggests a mechanistic relationship between castration and pro-tumorigenic immune signaling.

Finally, we tested the hypothesis that high translation rates were important to maintain AR-low prostate cancer heterogeneity using the *Pten*^*fl/fl*^*;4ebp1*^*M*^ mouse model. We discovered that 4EBP1^M^ induction severely depleted both basal and intermediate cells. Interestingly, hyper-proliferative basal cells were preferentially depleted, leading to a decrease in overall basal cell proliferation. In addition, EGFR and TNF signaling were decreased in the tumor microenvironment. We conclude that high translation rates are essential to maintain tumor heterogeneity in AR-low prostate cancer and may play a role in pro-tumorigenic cell-cell communication pathways. Based on these findings, we speculate that translation inhibitors may represent a therapeutic modality to decrease tumor heterogeneity. Overall, our work highlights multiple epithelial and immune cell types crucial to prostate cancer initiation and progression and elucidates interactions between specific cell populations that may facilitate castration resistance. Lastly, this work aims to provide a broad, searchable resource to the cancer research community. To this end, we have developed a publicly accessible and interactive website (available at https://atlas.fredhutch.org/hsieh-prostate/, username: hsieh, password: viz) that allows for cell- and gene-specific queries through all 50,780 cells analyzed in this study (Fig. S5).

## Materials & Methods

### Mice

PB-Cre mice were obtained from the Mouse Models of Human Cancer Consortium. Pten^fl/fl^ and Rosa-LSL-rtTA mice were obtained from the Jackson Laboratory. TetO-4ebp1^M^ mice were generated as previously described (Hsieh et al., 2015). All mice were maintained in the C57BL/6 background under specific pathogen–free conditions, and experiments conformed to the guidelines as approved by the Institutional Animal Care and Use Committee of Fred Hutchinson Cancer Research Center.

### Surgical castration and activation of the 4EBP1^M^ transgene

Surgical castrations were performed in 4- to 5-month-old mice under isoflurane anesthesia. Postoperatively, mice were monitored daily for 5 days. To activate the 4EBP1^M^ transgene, doxycycline (Sigma-Aldrich) was administered in the drinking water at 2 g/liter immediately after castration, and euthanasia was performed 8 weeks after castration.

### Tissue dissociation for single-cell RNA sequencing

Ventral prostate lobes from C57BL/6J WT, *Pten*^*fl/fl*^, *Pten*^*fl/fl*^*;4ebp1*^*M*^ mice were dissected, washed with chilled 1 X PBS, and then minced with a scalpel into small pieces (∼ 1 mm) in a petri dish. Paired lobes from a single mouse were collected and dissociated into one sample for scRNA-seq. The tissue was digested with DMEM/F12/Collagenase/Hyaluronidase/FBS (StemCell technologies, Vancouver, Canada) for 1 hours at 37°C on a slowly shaking/rotating platform. The tissue was further digested in 0.25% Trypsin-EDTA (Invitrogen, Carlsbad, CA) on ice for 30 minutes, and followed by suspension in Dispase (Invitrogen, Carlsbad, CA, 5 mg/mL) and DNase I (Roche Applied Science, Indianapolis, IN, 1 mg/mL). Any cell clumps were dissociated by gently pipetting up and down. The dissociated cells were then passed through 70 μm cell strainers (BD Biosciences, San Jose, CA) to achieve single cell suspension. The suspension was resuspended with 3 ml PBS (Life Technologies) with 2% fetal calf serum (FCS) (Gemini Bioproducts, West Sacramento, CA) and immediately placed on ice. Viable cells were counted by Vi-Cell XR Cell Viability Analyzer (Beckman Coulter, Brea, CA) and then diluted accordingly to reach the targeted cell concentration.

### Single-cell RNA sequencing library preparation

3’ single-cell RNA libraries were generated according to the protocol outlined in Single Cell 3’ Reagent Kits v2 User Guide 10X Genomics. Briefly, cells and reverse transcription reagents were partitioned into oil-based Gel Beads in Emulsion (GEMs), with each GEM containing a unique 10x barcode. Cells were then lysed and underwent reverse transcription resulting in barcoded cDNA. The cDNA was then collected and amplified prior to undergoing library construction in which P5, P7, and a unique sample index were added. The 10 libraries generated in this manner were pooled and sequencing was performed on an Illumina NovaSeq 6000 using the v1.5 S1-100 flow cell and reagent kit. Sequencing configuration was paired-end 26×8×96 and Illumina RTA version v3.3.5 was used. This generated a median of 58.7 million reads/sample with a median 54.60% saturation, median 90.70% Q30 fraction, 13,746 average reads/cell, and 6.2 reads/UMI.

### Alignment and Filtering of Reads

Two transgene transcripts (*Cre, rtTA-eGFP*) were added to the mm10 transcriptome to detect transgenes expressed in the *Pten*^*fl/fl*^ and *Pten*^*fl/fl*^*;4ebp1*^*M*^ mice. Kallisto v0.45.1 (Bray et al., 2016) was used to demultiplex all samples into FASTQ files and align reads to the modified mm10 transcriptome. DropletUtils package (Lun et al., 2019; Griffiths et al., 2018) was used to filter out empty or duplexed cell droplets. Cells with fewer than 200 or greater than 5000 detected genes, fewer than 500 or greater than 25,000 detected UMIs and cells with >15% mitochondrial reads were filtered from subsequent analysis.

### PCA, UMAP, and Clustering

R package Seurat v4.0.4 (https://satijalab.org/seurat/) was used to construct a Principal Component Analysis (PCA) for the entire dataset using the 2000 most variable genes as features. The Uniform Manifold Approximation and Projection (UMAP) dimension reduction technique was used for visualization and the R function “FindClusters()” with resolution = 0.2 was used to generate 43 clusters.

### Cell Type Identification

The SingleR package v1.6.1 was used to assign initial cell type identities to each cluster. These IDs were verified and refined using expression patterns of published biomarkers. For epithelial cells, cell subtypes (basal, intermediate, differentiated) were assigned using published gene signatures from other single-cell RNA sequencing projects. For immune cells, broad cell types (T cells, macrophages) were divided into activation states via known biomarkers (e.g. *Cd8a* for CD8 T cells and *Mrc1* for M2-activated macrophages). Stromal cell types were also determined via biomarkers.

### Relative Cell Abundance

To compare the abundance of specific cell populations while controlling for sample library size, the percentage composition of each sample was calculated by cell type. Statistical significance was generated via a negative binomial regression test to determine whether a given cell type was over- or under-represented between conditions.

### Gene Signature Enrichment

The GSVA package v1.40.1 (Hanzelmann et al., 2013) was used to generate composite scores for gene signatures such as a 20-gene AR activity signature or the 30-gene CCP proliferation signature. Due to the sparse nature of single-cell transcriptomes, the data was pseudo-bulked by sample and cell type to generate more robust analyses. Statistical analysis was performed via permutation test with 10,000 permutations.

### Differential Expression and Gene Set Enrichment Analysis

Differential gene expression was computed using Seurat functions with a threshold log2 fold-change > 0.25 or < -0.25 and FDR < 0.05. Upregulated and downregulated genes were further filtered by setting a log2 fold-change threshold = log2(1.25) = ∼0.32. Gene names were converted from mouse to human via the biomaRt package (Durinck et al., 2009) and GSEA was performed using the MsigDB database with the C2, C5, C6, C7, Hallmark, KEGG, BioCarta, and Reactome gene sets. Resulting enriched pathways were filtered via a threshold of FDR < 0.05.

### Trajectory, RNA Velocity, and Pseudotime

Monocle3 (Cao et al., 2019; Qiu et al., 2017; Trapnell et al., 2014) and velocyto (La Manno et al., 2018) were used to draw trajectory paths and RNA velocity maps, respectively, through the epithelial compartment of the *Pten*^*fl/fl*^ intact mice. Palantir (Setty et al., 2019) was used to delineate gene expression dynamics across pseudotime in basal and intermediate cells in *Pten*^*fl/fl*^ intact mice.

### Ligand-receptor Interactions

Ligand-receptor interactions between cell types were determined via the CellphoneDB package v2.0.0 (Efremova et al., 2020). Only interaction with p-value < 0.05 were included in the final analysis.

### Cell Cycle Assignment

Cell cycle phases for single cells was determined using the Seurat cell cycle function, which includes gene lists denoting the G2M and S phases. Gene names were converted from human to mouse using the biomaRt package to match our data, then the CellCycleScoring function was used to assign each cell either S, G2M, or G1 phase. Chi-squared test was used to determine whether the proportions of G1 cells was significantly different between clusters or conditions.

### Human Gene Signature of ADT Resistance and Correlation to Mouse Data

Tumor samples were laser capture microdissected from prostate cancer biopsies prior to undergoing six months of neoadjuvant androgen deprivation therapy plus enzalutamide and ranked based on volume of residual tumor in each patient, as previously described (Karzai et al., 2020; Wilkinson et al., 2021; Ku et al., 2021). Separately, differentially expressed genes (DEGs) derived from the PTEN null intact and castrate basal or intermediate cells were converted from mouse to human gene symbols using getLDS function from the biomaRt package v2.48.3 for R/Bioconductor (Durinck et al., 2009). Gene set enrichment analysis (GSEA) was performed on the basal vs. intermediate DEGs set against the top 50 genes associated with treatment resistance, and the top five leading edge genes from GSEA were used to stratify samples. Survival analysis was performed using the *survival* package in R on the TCGA prostate adenocarcinoma (The Cancer Genome Atlas Research Network, 2015) (n = 490) and MSKCC (Taylor et al., 2010) (n = 140) datasets. A cancer sample was considered “altered” if the expression of at least one of the five leading edge genes was greater than the 80th percentile for the entire cohort (TCGA or MSKCC, respectively).

### TCGA Analysis of TNF Activity

The Cancer Genome Atlas (TCGA) PRAD cohort containing 493 primary prostate tumor samples with RNA-seq expression values was utilized for analysis of signature scores. We used the RSEM values hosted by the cBioPortal (http://www.cbioportal.org, study: prad_tcga_pan_can_atlas_2018.) Single sample enrichment scores were calculated using GSVA (Hanzelmann et al., 2013) with default parameters using genome-wide log2 RSEM values as input. The pathways used were from MSigDBv7.4 (HALLMARK_TNFA_SIGNALING_VIA_NFKB) and the 10-gene androgen-regulated (AR) signature from Bluemn et al., 2017. In analyses restricted to samples with PTEN biallelic loss, 94 samples were used which had either 2 copy loss or 1 copy loss and a non-synonymous mutation annotated as a putative driver mutation in cBioPortal. Pearson’s correlation coefficient was used to study the relationships between signature scores shown in scatterplots using the cor.test function in R.

## Data and code availability

The code used to process and analyze the data is available at https://github.com/sonali-bioc/GermanosProstatescRNASeq/. All other data associated with this study are present in supplementary materials and tables.

## Interactive Website

The web-based data Atlas was developed utilizing open-source technologies, including React for the application framework, Material UI for interface components, and Apache EChart for visualizations. All data were extracted from Seurat HDF5 files into web-optimized CSV, Arrow, and Binary files. All site data and assets are stored in Amazon S3 and served through Amazon CloudFront, a global content delivery network (CDN) service built for high-speed, low-latency performance and security. The site is hosted at https://atlas.fredhutch.org/hsieh-prostate/ and can be accessed with username “hsieh” and password “viz.”

## Author contributions

A.A.G. and A.C.H. conceived of the study, performed the majority of analyses, and wrote and edited the manuscript. S.A. performed analyses and provided technical and coding assistance. A.C.H. and K.Y. performed the mouse dissection, A.L. and J.B. performed library preparation and single-cell RNA sequencing. E.G. and C.G. provided immunological expertise and assisted with the analysis of the cell-cell interaction studies. R.A. and R.G. provided technical guidance and assisted with data processing and analysis. Y.Z. performed statistical analysis and contributed code. M.S. provided trajectory and pseudotime expertise and assisted with Palantir analysis. P.S.N. provided advice on initial experimental design and I.C. and P.S.N. performed analyses of TCGA PRAD human data. A.T.K., S.W., and A.G.S. provided the human signature of ADT resistance and performed correlation analysis between human and mouse RNAseq data.

## Acknowledgments

We are grateful to the patients who participated in this study and their families. We thank members of the A.C.H. laboratory for helpful advice and discussions. We thank the Seattle RNA Metabolism group for critical discussion of the work. We thank L. Xin for sharing RNA-seq data and critical reading of the manuscript. This work was supported by NIH award R37 CA230617, the Pacific Northwest Prostate Cancer SPORE DRP (P50 CA097186), Burroughs Welcome Fund, Career Award for Medicine Scientists (1012314.02), and grants from the Emerson Collective (691630), and the Robert J. Kleberg Jr. and Helen C. Kleberg Foundation to A.C.H. A.A.G. received funding through an NIH T32 grant (T32CA080416) and E.T.G reviewed funding from the DoD BCRP Breakthrough Fellowship Award (W81XWH-19-1-0076). This research was also supported by P01 CA163227 and R01 CA234715 to P.S.N., the Prostate Cancer Foundation, the Genomics and Bioinformatics Shared Resource of the Fred Hutch/University of Washington Cancer Consortium (P30 CA015704) and the Scientific Computing Infrastructure at Fred Hutch funded by ORIP grant S10 OD028685.

## Figure Legends

**Fig. S1:**
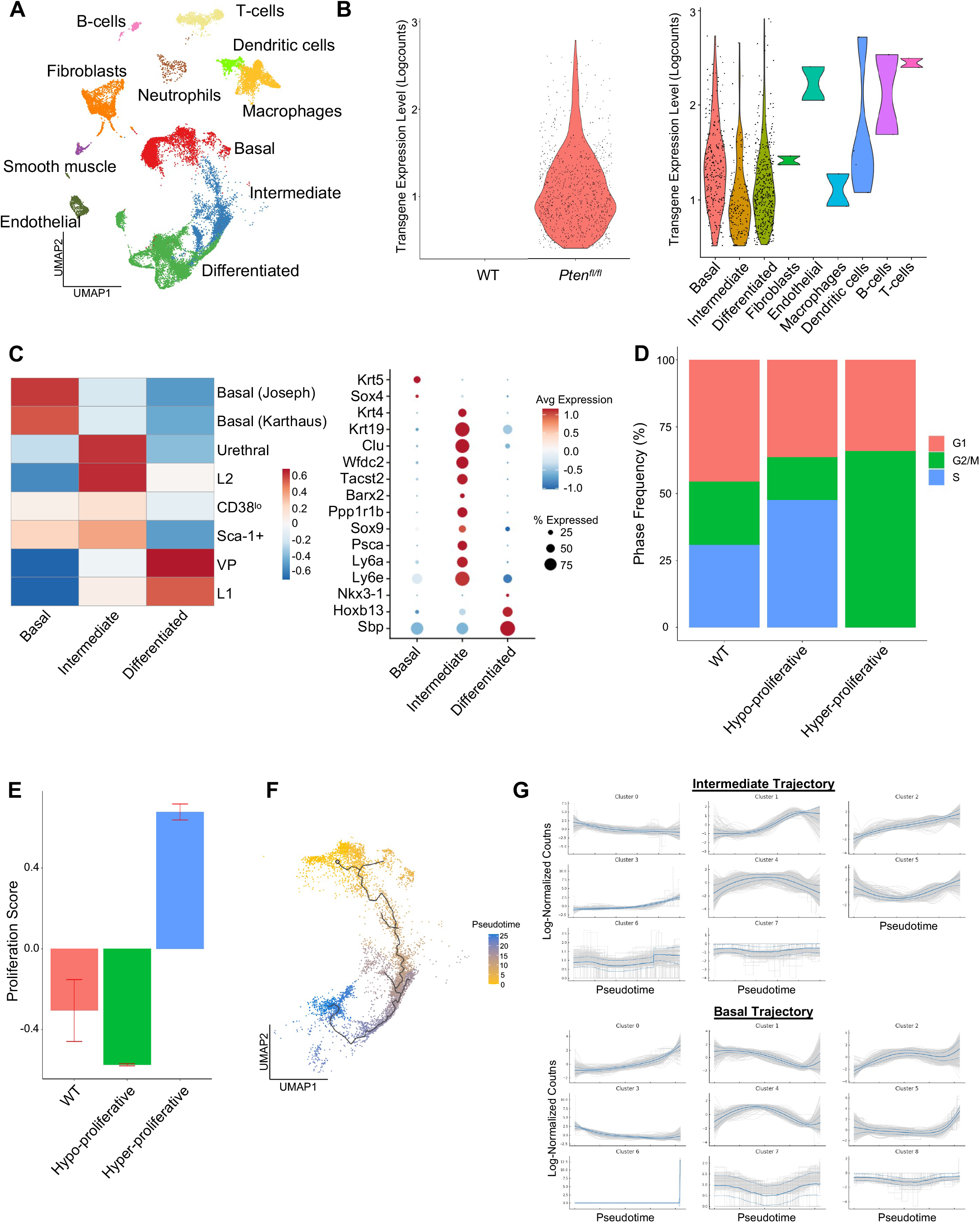
Epithelial Cells Contain Published Subtypes and Basal Proliferation is Subset-Specific. **A)** UMAP visualization of all cells in WT and *Pten*^*fl/fl*^ ventral prostates, colored and labeled by cell ID. **B)** Violin plots of *rtTA-eGFP* transgene expression. Left, transgene expression in WT and *Pten*^*fl/fl*^ mice. Right, expression in *Pten*^*fl/fl*^ cell types. **C)** Heatmap of composite scores of published prostate epithelial subtype signatures in basal, intermediate, and differentiated cells in WT mice (left). Dot plot of epithelial biomarker gene expression in WT mice (right). **D)** Bar plot of cell cycle phase assignments in WT basal cells and *Pten*^*fl/fl*^ hyper- and hypo-proliferative basal cells. **E)** Bar plot of CCP signature composite score in WT basal cells and *Pten*^*fl/fl*^ hyper- and hypo-proliferative basal cells (Data presented at +/- SD). **F)** Trajectory analysis of *Pten*^*fl/fl*^ epithelial cells. **G)** Top 3000 highly variable genes in *Pten*^*fl/fl*^ basal and intermediate cells, clustered by expression pattern along the basal-intermediate (top) or hypo-proliferative basal-hyper-proliferative basal (bottom) trajectories drawn via Palantir (Fig **1I**).

**Fig. S2:**
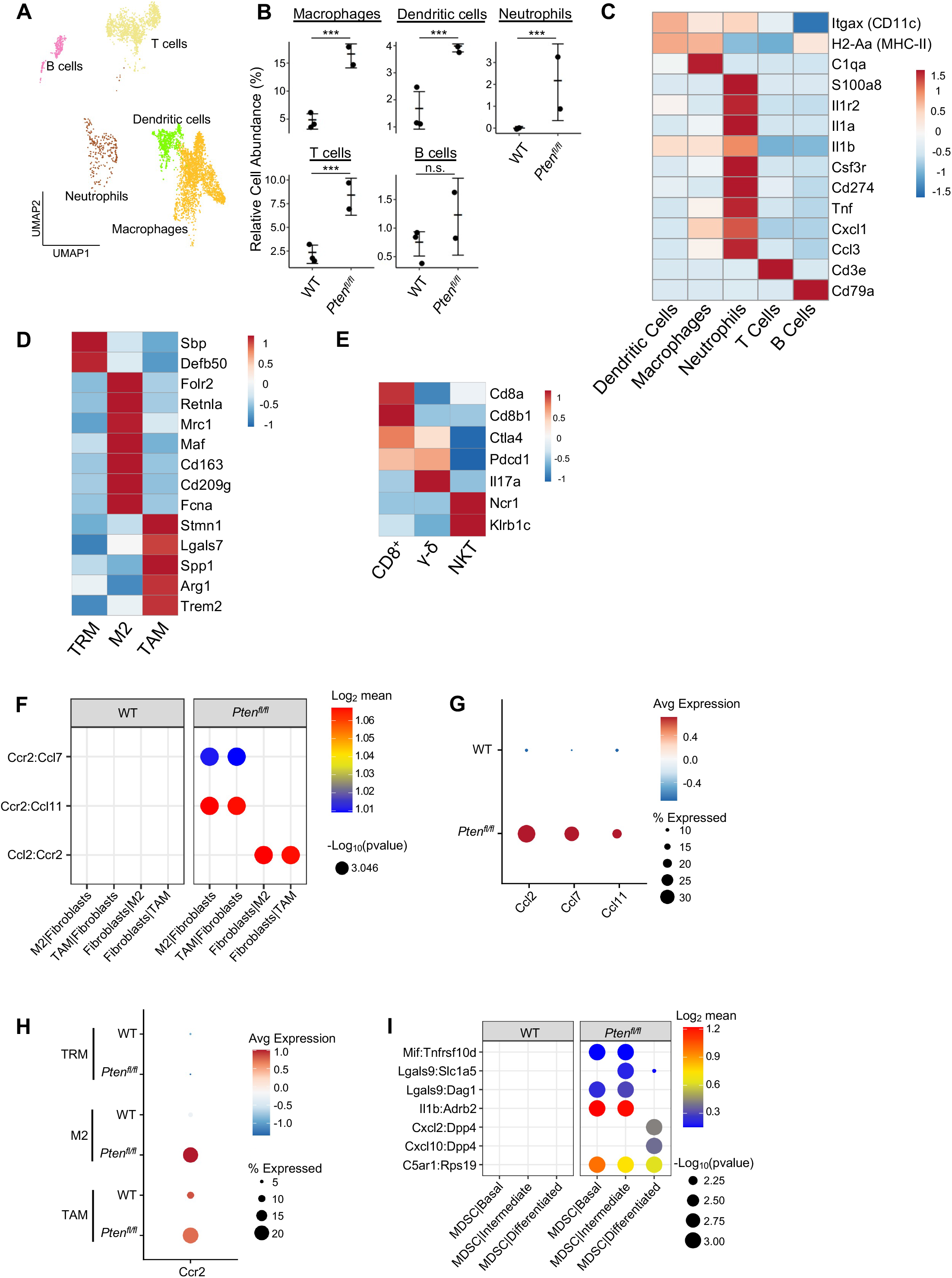
Immune Cells Contain Pro-Tumorigenic Subtypes and Macrophages are Recruited by Fibroblast Signaling. **A)** UMAP of immune cells in WT and *Pten*^*fl/fl*^ prostates, labeled by cell types. **B)** Relative abundance of immune cell types in WT and *Pten*^*fl/fl*^ mice (***p<0.001, negative binomial regression test). **C)** Heatmap of immune cell type biomarker expression in WT and *Pten*^*fl/fl*^ mice; neutrophil cell express MDSC markers. Log-transformed read counts. **D)** Heatmap of marker expression in macrophage cell subtypes. Log-transformed read counts. **E)** Heatmap of marker expression in T cell subtypes. Log-transformed read counts. **F)** Plot of ligand-receptor interactions between fibroblast and macrophage subtypes. **G)** Dot plot of *Ccr2* ligand expression in fibroblasts in WT and *Pten*^*fl/fl*^ ventral prostates. **H)** Dot plot of *Ccr2* expression in macrophage subtypes in WT and *Pten*^*fl/fl*^ ventral prostates. **I)** Dot plot of signaling interactions between epithelial cells and MDSCs. n.s. = not significant.

**Fig. S3:**
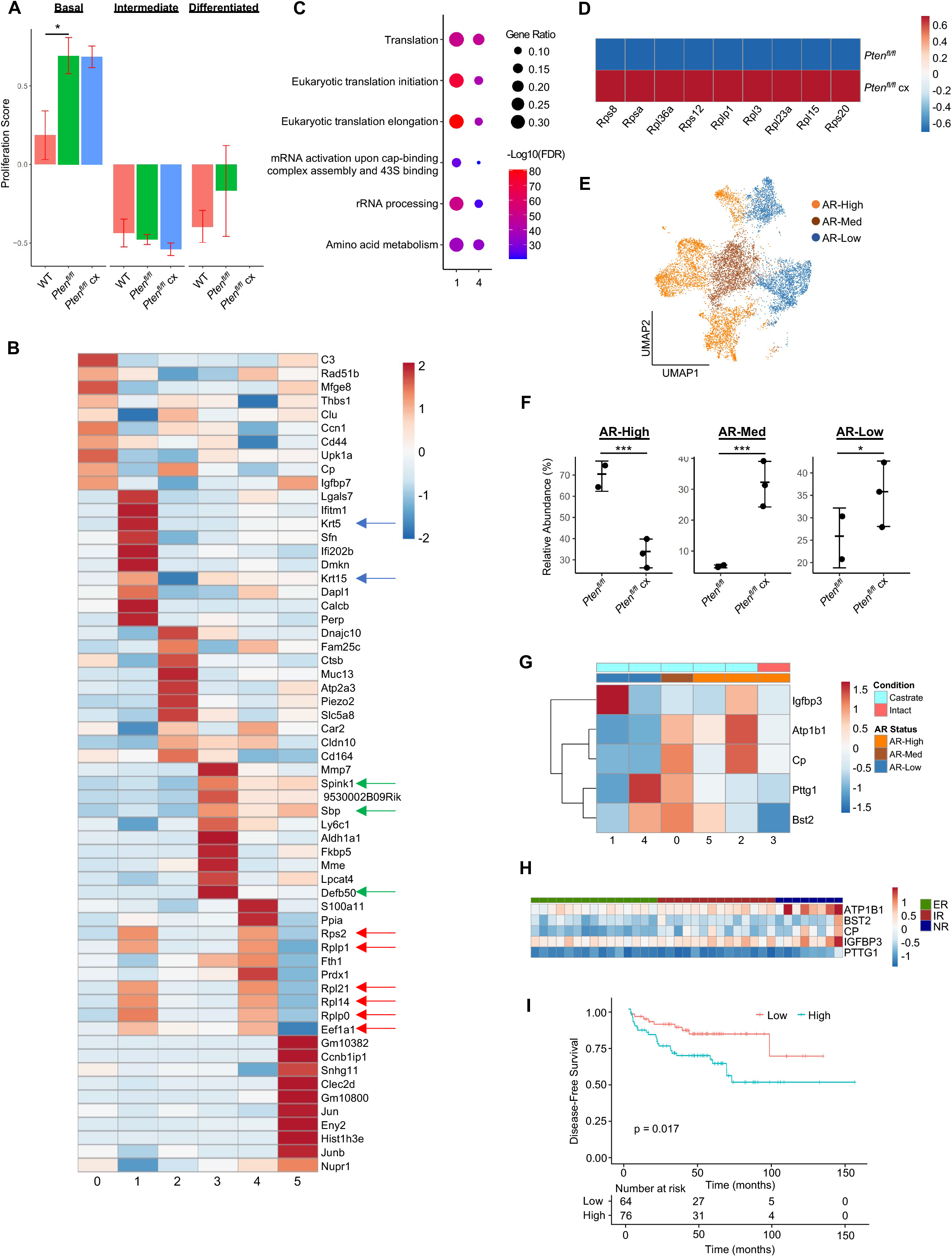
Castration-Resistant Intermediate Cells are Phenotypically Diverse and Correlate with Treatment Resistance in Human Patients. **A)** Composite score of CCP signature in WT, *Pten*^*fl/fl*^ intact, and *Pten*^*fl/fl*^ cx epithelial cells (Data presented at +/- SD, *p<0.05, permutation test) **B)** Heatmap of top differentially expressed genes across intermediate clusters 0-5. Blue arrows, basal markers; green arrows, AR-dependent genes; red arrows, ribosomal or translation machinery genes. **C)** Top GSEA results for genes upregulated in intermediate clusters 1 and 4. All pathways are enriched with FDR < 0.05. **D)** Heatmap of ribosomal gene expression in basal cells in *Pten*^*fl/fl*^ intact and *Pten*^*fl/fl*^ cx mice. **E)** UMAP visualization of AR signaling status in intermediate cells in *Pten*^*fl/fl*^ intact and *Pten*^*fl/fl*^ cx mice. **F)** Relative abundance of intermediate cells with high, medium, or low AR signaling in *Pten*^*fl/fl*^ intact and *Pten*^*fl/fl*^ cx mice (Data presented at +/- SD, *p<0.05, ***p<0.001, negative binomial regression test). **G)** Heatmap of top 5 resistance genes in intermediate clusters, labeled by AR status and condition (intact or castrate). **H)** Heatmap of the top 5 genes enriched in castrated intermediate cells (**Fig. 3I**) and their expression in ADT non-responder (NR), intermediate responder (IR), and excellent responder (ER) patients. **I)** Kaplan-Meier curve of disease-free survival of patients in the Taylor et al. (2010) cohort, separated by expression of the top 5 resistance genes in castrated intermediate cells. Red line, normal expression of top 5 genes; blue line, patients with expression of at least 1 gene with TPM in the 80^th^ percentile or above.

**Fig. S4:**
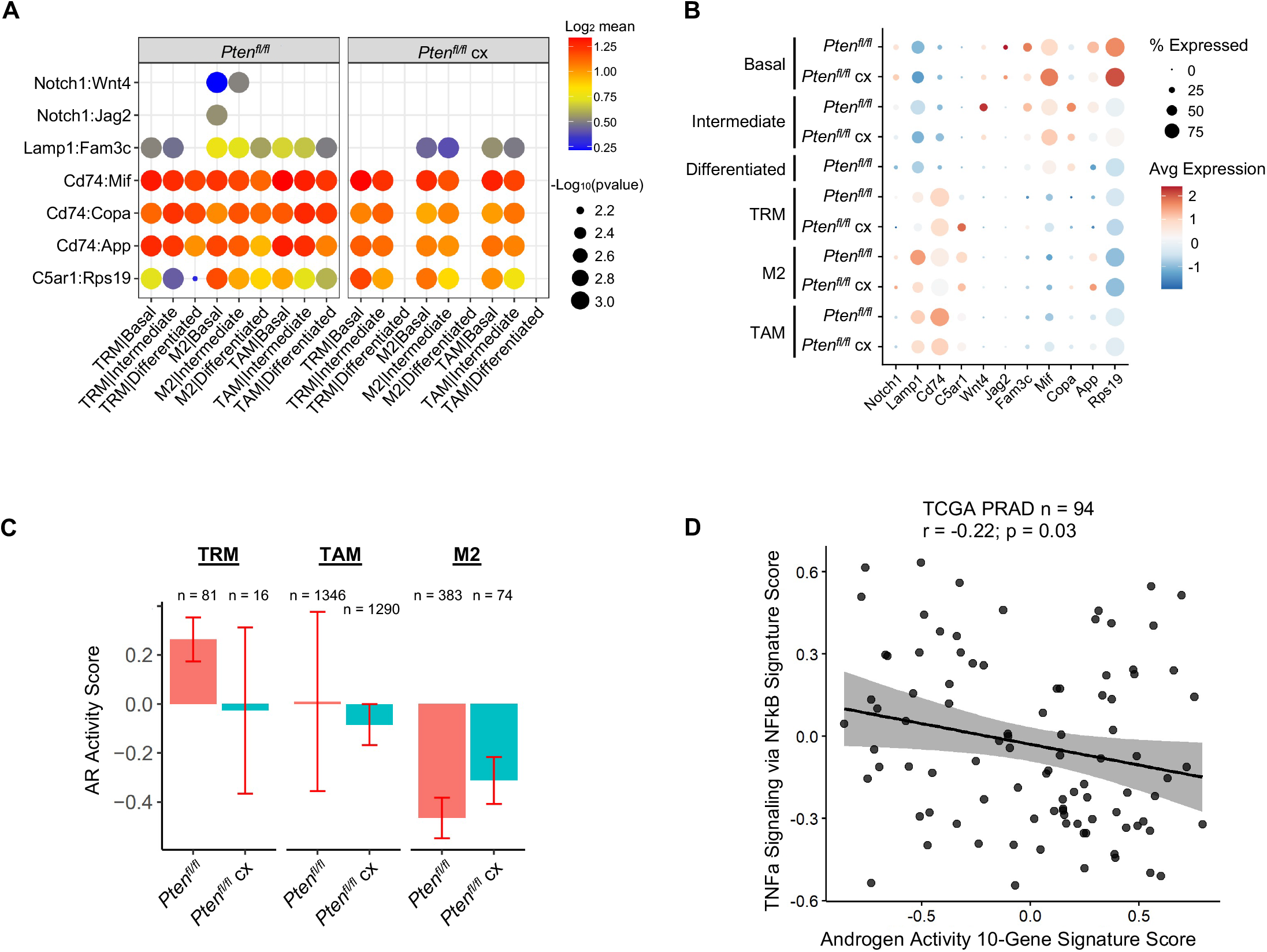
Epithelial-Mediated Macrophage Recruitment is not Interrupted by Castration. **A)** Plot of signaling interactions between macrophage subtypes and epithelial cells in *Pten*^*fl/fl*^ intact and *Pten*^*fl/fl*^ cx prostates. **B)** Dot plot of epithelial ligand and macrophage receptor gene expression in *Pten*^*fl/fl*^ intact and *Pten*^*fl/fl*^ cx ventral prostates. **C)** Composite score of AR signaling signature in macrophage subtypes in *Pten*^*fl/fl*^ intact and *Pten*^*fl/fl*^ cx prostates (Data presented at +/- SD). **D)** Scatter plot of TCGA PRAD study patient signature composite scores, filtered for patients harboring PTEN mutations. Y-axis, TNF signaling signature score; X-axis, AR signaling signature score (Pearson’s correlation).

**Fig. S5:**
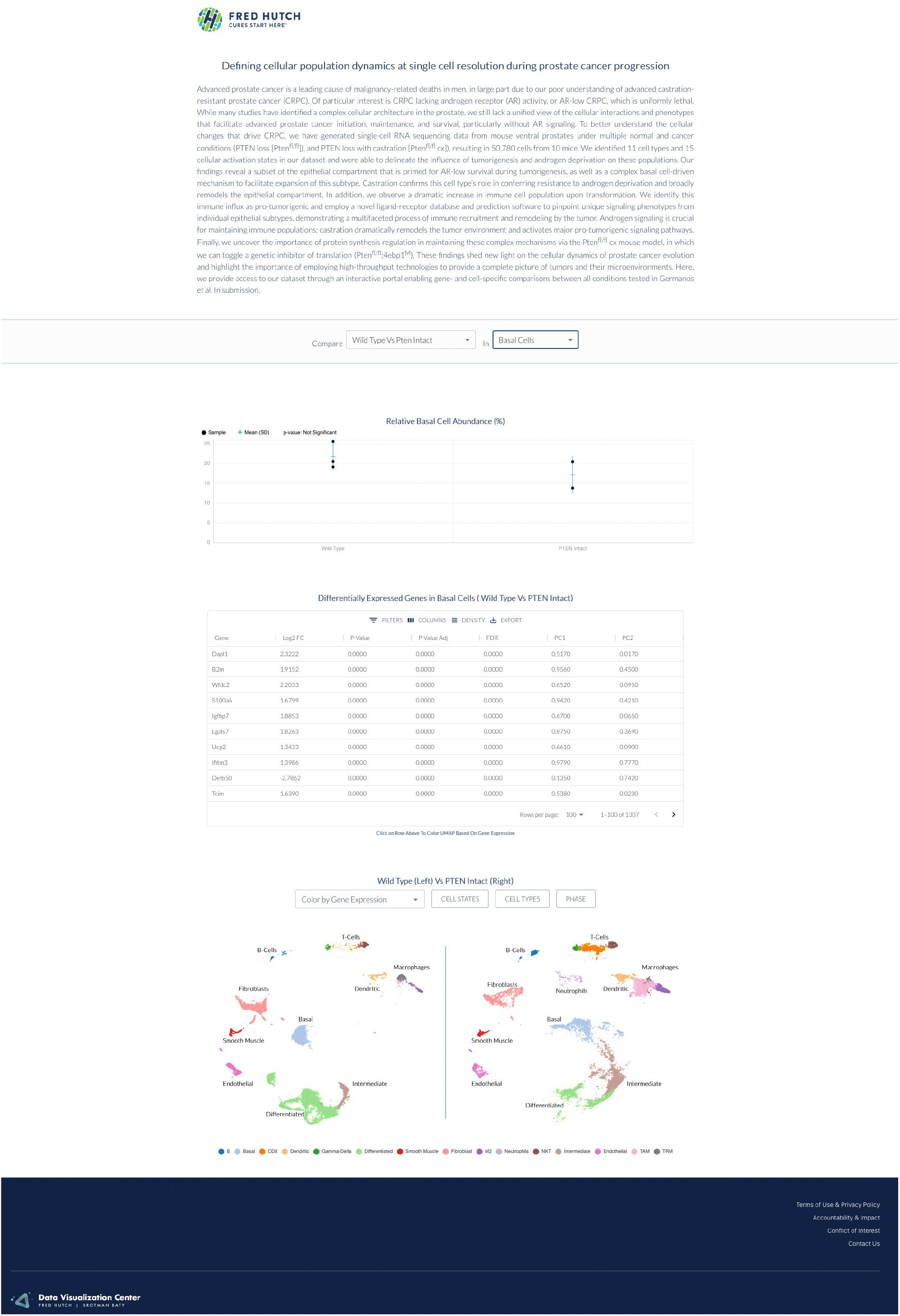
Sample Image of Interactive Portal, Enabling Gene- and Cell- Specific Comparisons Between Mouse Ventral Prostates.

